# Targeted Degradation of MYC2 through MPK6 Phosphorylation Reveals Mechanisms and Implications for Jasmonic Acid Signaling

**DOI:** 10.1101/2024.10.04.616682

**Authors:** Jong Hee Im, Seungmin Son, Man-Young Jung, Jae-Heung Ko, Kyung-Hwan Han

## Abstract

MYC2 is a key regulator in the Jasmonic acid (JA) signaling pathway, yet the mechanisms governing its stability remain unclear. In this study, we demonstrate that MYC2 is degraded by MPK6 under JA signaling. Through yeast two-hybrid and co-immunoprecipitation assays, and *in vitro* kinase assay, we show that MPK6 directly interacts with and phosphorylates MYC2. Additionally, MYC2 transcriptional activity is enhanced in the *mpk6* mutant. Further, MPK6 phosphorylates MYC2 at threonine 328, leading to its degradation. These findings suggest that MPK6 drives MYC2 degradation by specifically phosphorylating threonine 328 within the JA signaling pathway.

## Introduction

Jasmonic acid (JA) is a key phytohormone that regulates various plant processes, including growth, development, and stress responses. The regulation of JA signaling is mediated by several key components, including CORONATINE INSENSITIVE 1 (COI1), JASMONATE-ZIM-DOMAIN (JAZ) proteins, and MYELOCYTOMATOSIS 2 (MYC2) (Zander et al., 2020; Hewedy et al., 2023). Under conditions of inactive JA signaling, the activity of the basic helix-loop-helix (bHLH) transcription factor MYC2 is repressed by JAZ proteins, which form a corepressor complex with TOPLESS (TPL), HISTONE DEACETYLASE (HDA), and NOVEL INTERACTOR OF JAZ (NINJA) (Pauwels et al., 2010). To activate the JA signaling, JA-isoleucine conjugate (JA-Ile) induces the interaction between JAZ proteins and the Skp-Cullin-F-box (SCF) component COI1, leading to the ubiquitin-mediated degradation of JAZ via the 26S proteasome (Thines et al., 2007). This degradation relieves the repression on MYC2, allowing it to activate the expression of various JA-responsive genes. Therefore, deciphering the regulatory mechanisms of MYC2 is critical for advancing our understanding of plant responses to stress conditions and JA signaling.

Phosphorylation-based protein regulation serves as a key mechanism in numerous signaling pathways, playing a vital role in controlling protein activity and stability (Zhang et al., 2023). MITOGEN-ACTIVATED PROTEIN KINASE 6 (MPK6), an evolutionarily conserved serine and threonine kinase, is involved in various plant cellular signaling pathways, mediating the conversion of extracellular stimuli into intracellular responses (Zhang and Zhang, 2022). MPK6 is important in plant development and disease resistance (Sun and Zhang, 2022). Moreover, MPK6 phosphorylates and destabilizes the bHLH transcription factor INDUCER OF CBF EXPRESSION 1 (ICE1) conferring cold tolerance (Li et al., 2017). Additionally, MPK6 increases salt tolerance by degrading the cytokinin signaling component ARABIDOPSIS RESPONSE REGULATOR 1 (ARR1), ARR10, and ARR12 (Yan et al., 2021).

Given the role of MYC2 as a pivotal regulator in the JA signaling pathway, influencing a multitude of downstream genes (Zander et al., 2020), understanding the regulatory dynamics of MYC2 is essential for unraveling plant responses to stress conditions and hormonal signals.

Previous studies on the regulation of MYC2 by MPK6 have reported conflicting findings. Takahashi et al. suggested that MPK6 negatively regulates MYC2 (Takahashi et al., 2007) while Sethi et al. proposed a positive regulation (Sethi et al., 2014). However, the precise genetic and biochemical details of this interaction remain unclear.

In this study, we demonstrate that MPK6 directly phosphorylates MYC2 at threonine 328, leading to the targeted degradation of MYC2. This discovery provides important insights into the JA signaling pathway and its role in plant physiology, offering a clearer understanding of how MYC2 stability is controlled under JA signaling.

## Results

### Activated MPK6 directly phosphorylates and degrades MYC2 protein

In the previous study, the expression of *MYB46*, a direct target gene of MYC2, was significantly increased in *mpk6* plants (*mpk6-4*) (Kohoutová et al., 2015) under JA treatment conditions compared to Col-0 (Im et al., 2024), indicating the suppressive effect of MPK6 on the activity of MYC2. To further substantiate this finding, we evaluated the expression levels of two MYC2 direct target genes, *LIPOXYGENASE2* (*LOX2*) and *VEGETATIVE STORAGE PROTEIN1* (*VSP1*), in the plants with overexpression of *MYC2* in Col-0 (*MYC2OX*) or simultaneous overexpression of *MYC2* and *CONSTITUTIVELY ACTIVE MPK6* (*CAMPK6*) in Col-0 (*MYC2OX*/*CAMPK6OX*). Notably, the expression of these MYC2 targets was upregulated in the *MYC2OX* plants but significantly reduced in the *MYC2OX*/*CAMPK6OX* plants compared with *MYC2OX* (Fig. 1A), reinforcing the notion that MPK6 negatively impacts the transcriptional activity of MYC2.

**Figure 1.**
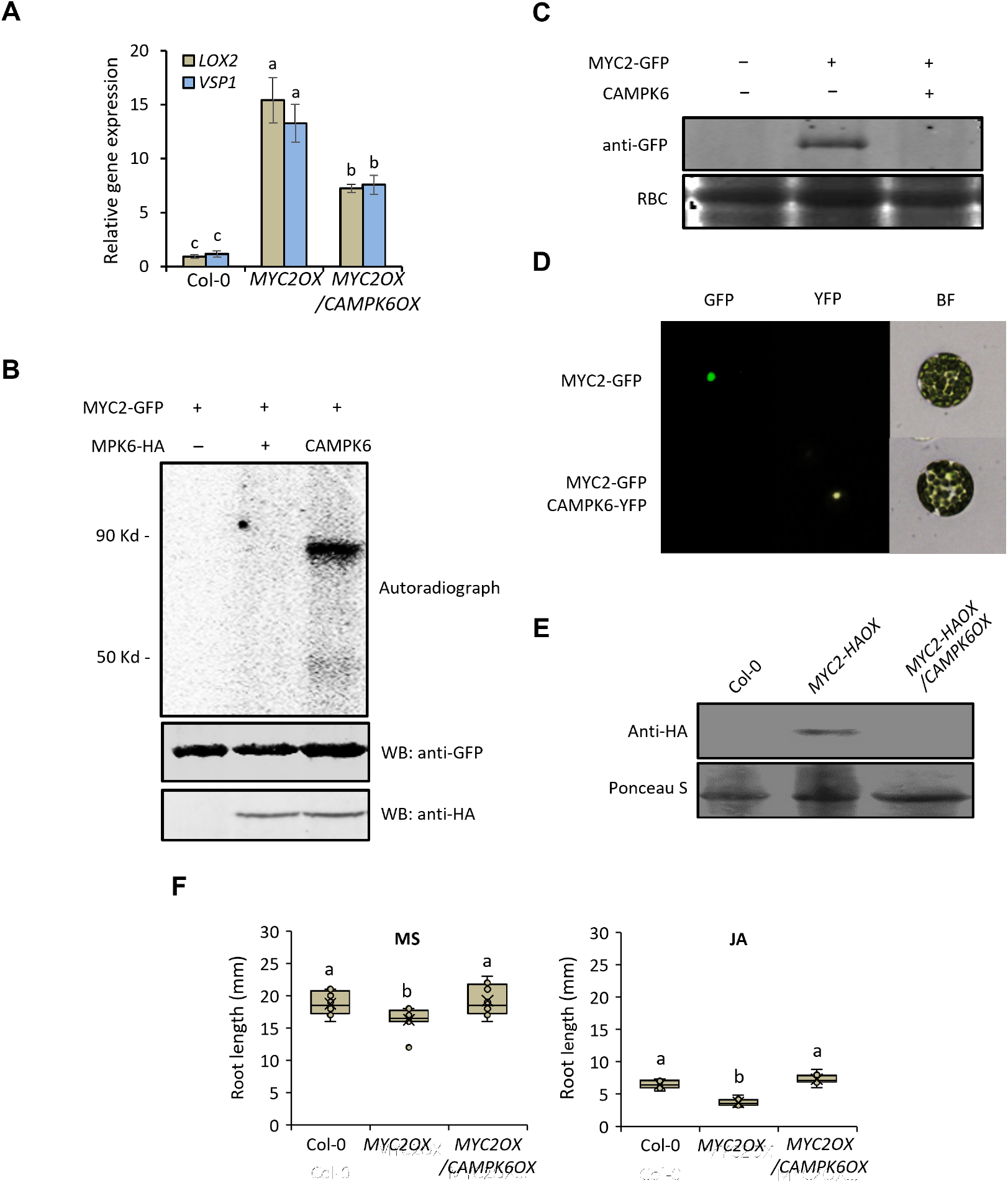
MPK6 negatively regulates MYC2 via phosphorylation-dependent protein degradation. **(A)** Gene expression analysis of *LOX2* and *VSP1* in Col-0, *MYC2OX* and *MYC2OX*/*CAMPK6OX* plants. Gene expressions were determined with RT-qPCR. Values are means of triplicate-independent biological repeats with SEs. Different letters indicate significant differences according to ANOVA (*P* < 0.05). **(B)** *In vitro* kinase assay of MYC2 with MPK6 and CAMPK6. MYC2-GFP, MPK6-HA, and CAMPK6-HA were respectively expressed in Arabidopsis leaf protoplasts (ALPs). After the expression, immunoprecipitations were carried out with anti-HA antibody and anti-GFP antibody, and then *in vitro* kinase assay was carried out. **(C)** Protein blot analysis of MYC2. C-terminal GFP conjugated MYC2 was expressed in ALPs with or without CAMPK6 co-expression. Anti-GFP antibody was used for detection of MYC2 protein and Ponceau S was used as loading control. **(D)** MYC2-GFP signal in CAMPK6 co-expression. C-terminal *GFP* conjugated *MYC2* was expressed in ALPs with or without C-terminal *YFP* conjugated *CAMPK6*. After the expression, the GFP and YFP signals were detected with a fluorescence microscope. BF, Bright field. **(E)** Comparing protein stability of MYC2-HA in *MYC2-HAOX* and *MYC2-HAOX*/*CAMPK6OX* plants. Total protein was isolated from 4-day-old Arabidopsis seedlings and carried out protein blot analysis with an anti-HA antibody. **(F)** Comparing primary root length in Col-0 and the transgenic plants under normal and JA conditions. Col-0, *MYC2OX*, and *MYC2OX*/*CAMPK6OX* seeds were germinated and grown for 4 days in solid MS medium with or without 50 uM MJA and taken the image of the primary root (*n* = 9). Values are means of biological repeats with SEs. Different letters indicate significant differences according to ANOVA (*P* < 0.05).

Building on the finding that MPK6 directly phosphorylates MYC2, we first investigated the nature of the interaction of the two proteins using a yeast two-hybrid assay. The results showed that CAMPK6 interacts with the 328 - 446 amino acid region of MYC2, which correlated with a potential MPK binding site (Eukaryotic Linear Motif, http://elm.eu.org/; Fig. S1). Subsequent co-immunoprecipitation experiments verified this interaction (Fig. S2). We used *in vitro* kinase assay to show that MYC2 is phosphorylated by CAMPK6 but not by MPK6 (Fig. 1B). Furthermore, we observed a notable reduction in MYC2 protein stability by CAMPK6 both in Arabidopsis (*Arabidopsis thaliana*) leaf protoplasts (Fig1. C and D) and *in planta* (Fig. 1E).

We then tested our hypothesis that MPK6-dependent MYC2 protein degradation correlates with physiological responses. The *MYC2OX* plants had shorter primary roots compared to Col-0 plants as previously shown (Jung et al., 2015). However, root length in the *MYC2OX*/*CAMPK6OX* plants was restored to that of the Col-0 with or without JA treatments (Fig. 1F and Fig. S3), suggesting a functional interplay between MYC2 and MPK6 that influences root growth. These results suggest that active MPK6 directly interacts with and phosphorylates MYC2, which in turn leads to the degradation of MYC2.

### MYC2 activity is increased in *mpk6* mutant

To confirm the effect of MPK6 on MYC2 transcriptional activity, we analyzed the MYC2 activity in the *mpk6* mutant. The gene expression pattern of *LOX2* was similarly observed with *MYB46* (Fig. 2A) (Im et al., 2024). Further, the expression of *LOX2* was increased in the *MYC2OX* plants but the increase was significantly higher in the *MYC2OX*/*mpk6* plants, while the level of trans-*MYC2* transcription is similar in both the transgenic plants (Fig. 2B and Fig. S4). These findings collectively underscore that MPK6 acts as a negative regulator of MYC2.

**Figure 2.**
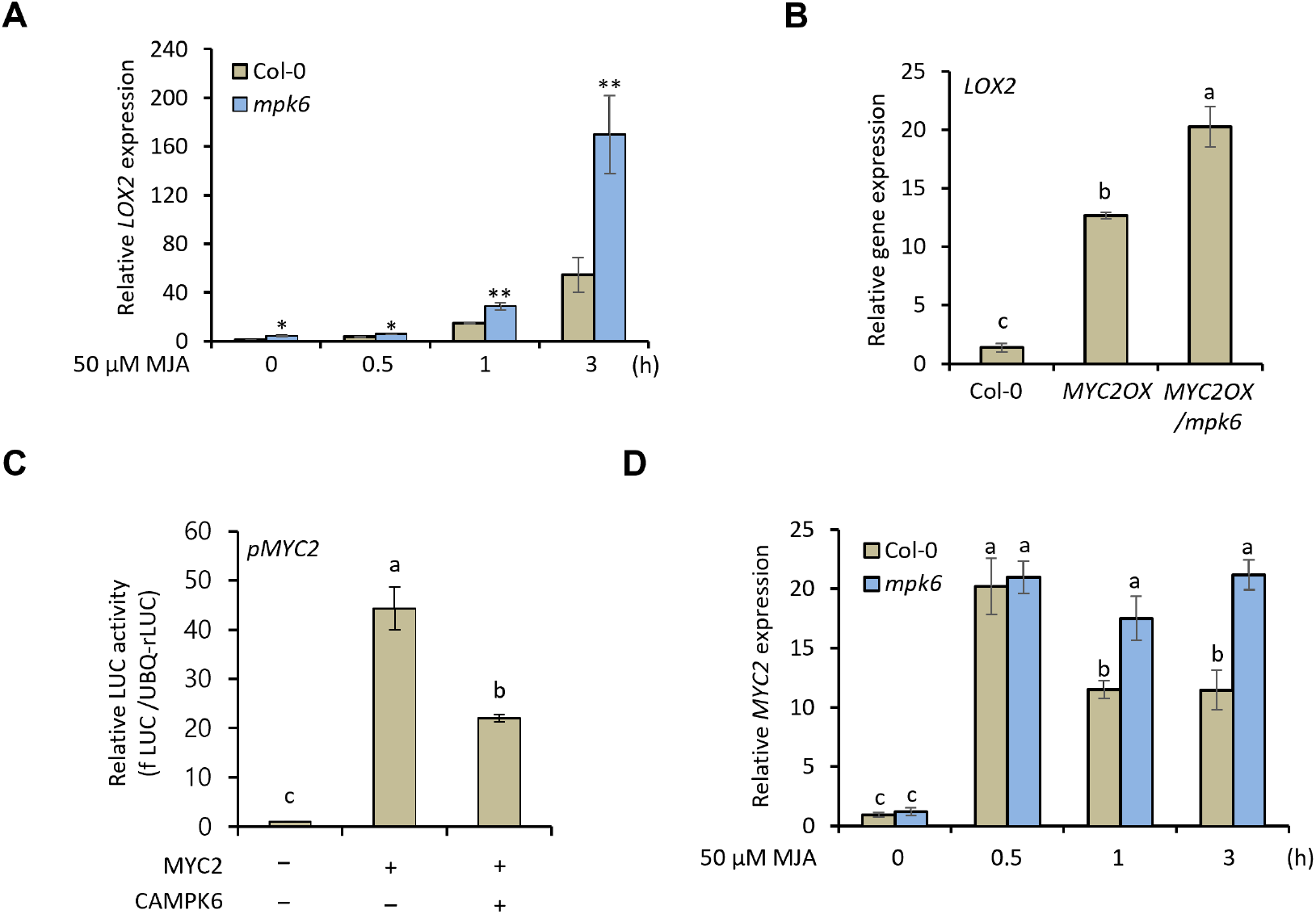
Transcriptional activity of MYC2 in *mpk6*. **(A)** Gene expression of *LOX2* in Col-0 and *mpk6* under JA treatment. 4-day-old Col-0 and *mpk6* seedlings were treated with 50 uM MJA for designated time points. Total RNA was extracted from the seedlings and cDNA was synthesized with it. Gene expressions were determined with RT-qPCR. Values are means of triplicate-independent biological repeats with SEs (** *P* < 0.01, and * *P* < 0.05). **(B)** Gene expression of *LOX2* in Col-0, *MYC2OX* and *MYC2OX*/*mpk6*. Total RNA was isolated from 4-day-old seedlings and gene expressions were determined with RT-qPCR. Values are means of triplicate-independent biological repeats with SEs. Different letters indicate significant differences according to ANOVA (*P* < 0.05). **(C)** Promoter transient assay of *MYC2* with MYC2 effector protein and CAMPK6. C-terminal *fLUC* conjugated *MYC2* promoter was expressed in ALPs with or without MYC2, and CAMPK6 co-expressions as designated combinations. *UBQ*::*rLUC* was used as an expression control. Values are means of triplicate-independent biological repeats with SEs. Different letters indicate significant differences according to ANOVA (*P* < 0.05). **(D)** Gene expression analysis of *MYC2* in Col-0 and *mpk6* under JA treatment. 4-day-old Col-0 and *mpk6* seedlings were treated with 50 uM MJA until 3 h in a time-dependent manner and carried out RT-qPCR for analysis of *MYC2* expression. Values are means of triplicate-independent biological repeats with SEs. Different letters indicate significant differences according to ANOVA (*P* < 0.05).

Given that MYC2 exhibits self-transcriptional activity (Wang et al., 2019) which was notably diminished by CAMPK6 (Fig. 2C), we evaluated *MYC2* expression level in both Col-0 and *mpk6* mutants following JA treatment. An increase in *MYC2* expression was observed at 0.5 h after the application of 50 μM JA, followed by a decline within 1 h in Col-0. However, this decline was not evident in the *mpk6* mutants (Fig. 2D). These findings collectively underscore that endogenous MPK6 acts as a negative regulator of MYC2 transcriptional activity in the context of JA signaling.

### MPK6 phosphorylates threonine 328 of MYC2

To pinpoint the exact phosphorylation target site of MYC2 by MPK6, we conducted *in vitro* kinase assays with CAMPK6, MYC2, and two phospho-mutants of two putative MPK6 target phosphorylation sites (S123A, T328A) of MYC2 (Zhai et al., 2013; Sethi et al., 2014). Here, we show that T328 is the phosphorylation target site by MPK6. In the kinase assay with two phospho-mutants, the phosphorylation of MYC2^T328A^ by CAMPK6 was notably less than that of both wild-type MYC2 and MYC2^S123A^ (Fig. 3A). To confirm this, we developed anti-MYC2^pT328^ antibody (ENPNLDP_P_TPSPVHSQT) and confirmed the specificity in Arabidopsis protoplasts (Fig. S5). In WT Col-0 plants, T328 of MYC2 was strongly phosphorylated at 10 min and then reduced at 20 min after 50 μM JA treatment but the phosphorylation was not detected in the *mpk6* mutant (Fig. 3B). Furthermore, MYC2 ^T328A^ more rapidly moved than WT MYC2 in co-expression with CAMPK6 (Fig. 3C).

**Figure 3.**
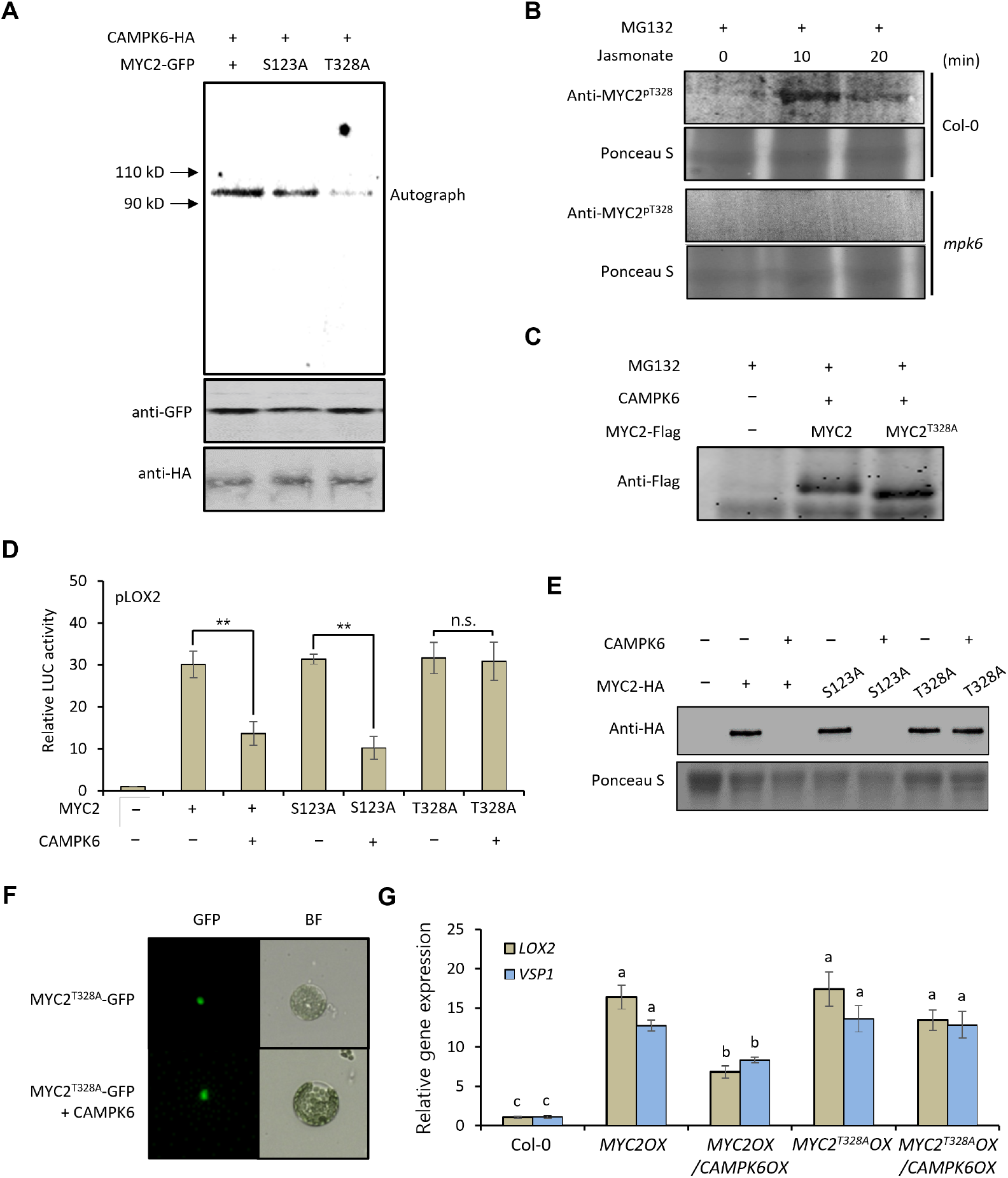
MPK6 degrades MYC2 through threonine 328 phosphorylation. **(A)** *In vitro* kinase assay of CAMPK6 with MYC2 and two phosphomutants. **(B)** Protein blot analysis of MYC2^pT328^ in JA treatment. Three-day-old Col-0 seedlings were treated with 10 μM MG132 for 24h and additionally treated with 50 μM Methyl JA for designated time points. Total protein was isolated from the seedlings and carried out protein blot analysis with an anti-MYC2^pT328^ antibody (ENPNLDPT_P_PSPVHSQT). **(C)** Gel shift assay of MYC2. *MYC2-Flag* and *MYC2*^*T328A*^*-Flag* were respectively expressed in 1 mL APLs with *CAMPK6*. After 1 h incubation, 1 μl of 10 mM MG132 was added and incubated additionally for 9 h. Total protein was isolated from the cells and separated with 10 % polyacrylamide gel with SDS-PAGE system for 2.5 h, subsequently carried out protein blot analysis with anti-Flag antibody. **(D)** Promoter transient assay of *LOX2* with MYC2 or its phosphomutants. C-terminal *fLUC* conjugated *LOX2* promoter was expressed in APLs with or without MYC2 and CAMPK6 as designated combinations. *UBQ-rLUC* was used as an expression control. Values are means of triplicate-independent biological repeats with SEs. Different letters indicate significant differences according to ANOVA (*P* < 0.05). **(E)** Protein blot analysis of MYC2-HA and its phosphomutants in the presence of CAMPK6 co-expression. **(F)** MYC2^T328A^-GFP signals. C-terminal *GFP* conjugated *MYC2*^*T328A*^ was expressed in ALPs with or without *CAMPK6* co-expression. GFP signal was detected with a fluorescence microscope. **(G)** Gene expressions of *LOX2* and *VSP1* in Col-0, *MYC2OX, MYC2OX*/*CAMPK6OX, MYC2*^*T328A*^*OX*, and *MYC2*^*T328A*^*OX*/*CAMPK6OX* plants. Total RNA was isolated from 4-day-old seedlings. Gene expressions were determined with RT-qPCR. Values are means of triplicate-independent biological repeats with SEs. Different letters indicate significant differences according to ANOVA (*P* < 0.05).

To identify the biological meaning of this, we analyzed the transcriptional activity of MYC2 and its phoshomutants with or without CAMPK6. The MYC2-mediated activity of the *LOX2* promoter was decreased in the presence of CAMPK6 in both MYC2 and MYC2^S123A^ but not in MYC2^T328A^ (Fig. 3D). This observation was further supported by the analysis of the stability of MYC2 and its phospho-mutants in the context of CAMPK6 co-expression, which showed that both MYC2 and MYC2^S123A^ proteins were destabilized with CAMPK6 co-expression, whereas MYC2^T328A^ remained stable (Fig. 3E and F). This was consistent with the *in planta* experiments, where *LOX2* and *VSP1* expression levels were increased in the *MYC2OX* plants but were significantly diminished in the *MYC2OX*/*CAMPK6OX* plants (Fig. 1h). However, the expression levels of *LOX2* and *VSP1* in the *MYC2*^*T328A*^ overexpressing plants (*MYC2*^*T328A*^*OX*) were comparable to those in the *MYC2OX* plants and not decreased in the presence of *CAMPK6* (*MYC2*^*T328A*^*OX*/*CAMPK6OX*), unlike in the *MYC2OX*/*CAMPK6OX* plants. These findings confirm that MPK6 targets T328 on MYC2 for phosphorylation, leading to the phosphorylation-dependent degradation of the MYC2 protein.

## Discussion

MYC2 is a master regulator of JA signaling but the protein stability of MYC2 in early time of JA signaling is still elusive. The gene expression of *MYC2* is significantly increased after 0.5 h after 100 μM Methyl JA (MJA) treatment but the protein level of MYC2 and MYC2 target gene *LOX2* expression not much changed at that time point (Zhai et al., 2013), suggesting that MYC2 protein is negatively regulated by MJA treatment. Moreover, our previous study showed upregulation of the MYC2 target gene *MYB46* in *mpk6* mutant (Im et al., 2024). Based on these two pieces of evidence, we hypothesized that MPK6 negatively regulates MYC2.

To prove the hypothesis, we used CAMPK6 rather than wild-type MPK6 because MPK6 is not activated under normal conditions (Xu et al., 2014) and the specificity of CAMPK6 on the MPK6 substrate is not different (Fig. S1) (Berriri et al., 2012). CAMPK6 strongly phosphorylated MYC2 but not MPK6 (Fig. 1B). CAMPK6 reduced MYC2 target gene expression, degraded MYC2 protein, and recovered short primary root growth by MYC2 overexpression (Fig. 1), further confirming the negative regulation of MYC2 by MPK6 in *mpk6* mutant (Fig. 2).

Threonin 328 of MYC2 is important for MYC2 protein stability but the kinase that phosphorylates this site has not yet been identified (Zhai et al., 2013). Sethi et al. identified S123 as the target phosphorylation site by MPK6 but the data was not confirmed with genetical and biological pieces of evidence (Sethi et al., 2014). Based on these two reports, we generated non-phoshorable MYC2 mutant MYC2^S123A^ and MYC2^T328A^ and analyzed the phosphorylation by CAMPK6. Fig. 3 shows that CAMPK6 phosphorylates Threonin 328 of MYC2 and the phosphorylation is important for MYC2 protein stability.

Phosphorylation-dependent negative regulation of MYC2 by MPK6 has been suggested previously. MYC2 binding on the *MPK6* promoter is blocked by MPK6-dependent MYC2 phosphorylation (Ojha et al., 2023). In salt stress, MPK6 phosphorylates MYC2, and the MYC2 reduced promoter binding activity on the *P5CS1* promoter (Verma et al., 2020).

However, the opposite response was also reported with different phosphorylation sites. Blue light-dependent MYC2 S123 phosphorylation by MPK6 positively regulates MYC2 transcriptional activity (Sethi et al., 2014). This opposite response is explainable. Human ERK2 phosphorylates Elk-1 at the different phosphorylation sites in a time-dependent manner (Mylona et al., 2016), showing different phosphorylation could be differently regulated even with the same kinase phosphorylation. However, the detailed mechanism of the MPK6-mediated MYC2 regulation on this concept is still elusive.

In this study, we provide an insight into the complex molecular interplay between MPK6 and MYC2 in the JA signaling pathway and its impact on plant growth by pinpointing the exact phosphorylation target site on MYC2 protein. The identification of T328 as the pivotal phosphorylation site that influences MYC2 stability marks a critical advance in our understanding of plant responses to phytohormone and environmental changes, opening new possibilities for biotechnological innovations of crops.

## Materials and Methods

### Plant growth conditions and JA treatment

*A. thaliana* Col-0 seeds were germinated and grown in ½ liquid or solid MS medium contained 1% sucrose at 24°C. Methyl JA treatment for root length analysis, *Arabidopsis* seeds were germinated and grown in solid medium (1/2 MS plus 0.5% sucrose contained 50 mM Methyl JA [Methyl 3-oxo-2-(2-pentenyl) cyclopentaneacetate, 3-Oxo-2-(2-pentenyl) cyclopentaneacetic acid, methyl ester]) for 4 days and measured primary root length. Methyl JA (MJA) treatment for RT-qPCR, liquid MS-grown 4-day-old seedlings were subjected to 50 μM MJA for the desired time and harvested. MJA for protein blot analysis, 10 μM MG132 was treated on a three-day-old Arabidopsis seedling for 24 h, and treated 50 μM MJA for the desired time point. For protoplast generation, plants were grown in soil for 23 to 25 days under a photoperiod of 13 h light/11 h dark at 23°C. The humidity was adjusted to 50%.

### Plasmid generation

*MYC2*^*T328A*^ and *MYC2*^*T123A*^ were generated by site-direct mutagenesis (Im et al., 2014) and transformed into Col-0 and *CAMPK6OX* plants (Im et al., 2024). C-terminal *firefly luciferase* conjugated *MYC2* and *LOX2* promoters were used in a previous study (Son et al., 2021). *MYC2* was cloned into C-terminal *GFP* conjugated HBT vector for detection localization and protein blot analysis.

For protein blot analysis in protoplasts, MYC2, *MYC2*^*T328A*^ and *MYC2*^*T123A*^ were cloned into C-terminal *Flag* conjugated HBT vector (Im et al., 2014).

### Protein blot analysis

In seedlings, total protein was isolated from 4-day-old seedlings using protein extraction buffer [50 mM Tris (pH 7.4), 1% NP-40, 150 mM NaCl, 1 mM EDTA with protease inhibitor cocktail]. The concentration of total protein was measured with the Bradford assay (Bradford, 1976). In the protoplast, MYC2 and its phosphomutants were respectively expressed in protoplasts for 10 h with or without additional treatment or co-expression of CAMPK6. After the total protein extraction, the protein was separated with the SDS-PAGE system. After the protein was transferred to the PVDF membrane, an anti-HA antibody, anti-GFP antibody, anti-Flag (Thermo Fisher Scientific, USA), and anti-MYC2^pT328^ antibody (Abclone, Korea) were used as the primary antibody, and an HRP conjugated antibody was used as a secondary antibody.

### Generation of *Arabidopsis* mesophyll protoplasts and transient promoter assay

Protoplast generation and transient promoter assay were performed as described previously (Son et al., 2023). For a generation of effector proteins, *MYC2* and *CAMPK6* cDNA were cloned in *HA* or *YFP-tagged* HBT vector. Plasmid DNA was prepared using a Plasmid Plus Maxi kit (QIAGEN, USA), and used for transfection. 8 μg of *fLUC* conjugated MYC2 target promoter was co-expressed with 6 μg of *UBQ10-rLUC* plasmid. Then, added 34 μg of effector plasmid or empty vector for control, then transfected to 200 μl of protoplasts, and incubated for 6 h. For the reporter assay, The promoter activities were presented as the firefly luciferase/renilla luciferase ratio and normalized to the values obtained without treatment or effector expression.

For detection of the fluorescence signal, *MYC2-GFP* or *MYC2*^*T328A*^*-GFP* were transfected to Arabidopsis protoplasts with or without *CAMPK6-YFP*. After 10 h incubation, the fluorescence signals were detected with a fluorescence microscope (339 Leica DRAMA2).

### Gene expression analysis

Total RNA was extracted from 4-day-old *Arabidopsis* seedlings using RNeasy Plant Mini Kit (Qiagen). For cDNA synthesis, SuperScript™ II Reverse Transcriptase (Invitrogen) was used. qPCR was carried out with specific primers on 7500 Real-Time PCR System (Applied Biosystems) using comparative Ct method with Fast SYBR™ Green Master Mix (Applied Biosystems). *Actin* (Im et al., 2021) was used as an expression control.

### *In vitro* kinase assay

C-terminal *GFP* conjugated *MYC2* and C-terminal *HA* conjugated *MPK6-HA* or *CAMPK6* were respectively expressed in protoplasts and incubated for 10 h. The cells were lysed completely with kinase-lysis buffer (20 mM Tris [pH 7.5], 40 mM MgCl_2_, 5 mM EDTA, 1 mM DTT, 10 mM NaF, 10 mM Na_3_Vo_4_, 1× protease inhibitor cocktail, and 1% Triton X-100). For immunoprecipitation (IP) analysis, the protein extracts were incubated with 1.5 μL tag-specific antibodies at 4°C for 3 h and an additional 3 h after adding 10 μL A-agarose beads. The beads were washed with IP buffer (50 mM Tris [pH 7.5], 150 mM NaCl, 5 mM EDTA, 1 mM DTT, and 1× protease inhibitor cocktail) and then twice with kinase buffer (20 mM Tris [pH 7.5], 40 mM MgCl_2_, 5 mM EDTA, and 1 mM DTT). The kinase reaction was performed with radioactive ATP as described previously (Im and Yoo, 2014).

### Gel shift assay

*MYC2-Flag* and *MYC2*^*T328A*^*-Flag* were transfected with CAMPK6 and treated with 10 μM MG132. After an additional 10 h incubation, total protein was isolated from the samples, and the protein was separated with an SDS-PAGE system using 100 voltage for 2.5 h running time, then carried out protein blot analysis following the abovementioned method.

### Statistical analysis

All experiments were independently conducted at least three times, and the data were analyzed by *t*-test or ANOVA. Asterisks denote significant differences (* *P* < 0.05, ** *P* < 0.01) and different letters indicate statistical differences (*P* < 0.05).

## Acknowledgments

This research was supported by DOE Great Lakes Bioenergy Research Center (DOE BER Office of Science DE-SC0018409), the Brain Pool Program by the Ministry of Science and ICT through the National Research Foundation of Korea (RS-2023-00262576), Regional Innovation Strategy (RIS) through the National Research Foundation of Korea (NRF) (2023RIS-009) and National Research Foundation of Korea (NRF-2023R1A2C1004193).

## Author contributions

JHI and SS conceptualized the study and performed the experiments. J-HK and K-HH supervised the study. JHI, SS, J-HK, M-YJ, and K-HH wrote the manuscript. All authors read and approved the manuscript.

## Data availability

The data that support the findings of this study are openly available from the corresponding author upon reasonable request.

## Supporting Information

**Fig. S1** Yeast two-hybrid assay shows that MYC2 directly interacts with CAMPK6.

**Fig. S2** Co-immunoprecipitation assay and *in vitro* kinase assay show that MYC2 is directly phosphorylated by CAMPK6.

**Fig. S3** Primary root length of Col-0, *MYC2OX*, and *MYC2OX/CAMPK6OX* in MS with or without MJA.

**Fig. S4** Eexpression of *MYC2* in Col-0, *MYC2OX*, and *MYC2OX/mpk6*.

**Fig. S5** Test of specificity of anti-MYC2^pT328A^ antibody.

**Table S1** The primers used in this study.

